# NRV: An open framework for in silico evaluation of peripheral nerve electrical stimulation strategies

**DOI:** 10.1101/2024.01.15.575628

**Authors:** Thomas Couppey, Louis Regnacq, Roland Giraud, Olivier Romain, Yannick Bornat, Florian Kölbl

## Abstract

Electrical stimulation of peripheral nerves has been used in various pathological contexts for rehabilitation purposes or to alleviate the symptoms of neuropathologies, thus improving the overall quality of life of patients. However, the development of novel therapeutic strategies is still a challenging issue requiring extensive *in vivo* experimental campaigns and technical development. To facilitate the design of new stimulation strategies, we provide a fully open source and self-contained software framework for the *in silico* evaluation of peripheral nerve electrical stimulation. Our modeling approach, developed in the popular and well-established Python language, uses an object-oriented paradigm to map the physiological and electrical context. The framework is designed to facilitate multi-scale analysis, from single fiber stimulation to whole multifascicular nerves. It also allows the simulation of complex strategies such as multiple electrode combinations and waveforms ranging from conventional biphasic pulses to more complex modulated kHz stimuli. In addition, we provide automated support for stimulation strategy optimization and handle the computational backend transparently to the user. Our framework has been extensively tested and validated with several existing results in the literature.

**Author summary:** Electrical stimulation of the peripheral nervous system is a powerful therapeutic approach for treating and alleviating patients suffering from a large variety of disorders, including loss of motor control or loss of sensation. Electrical stimulation works by connecting the neural target to a neurostimulator through an electrode that delivers a stimulus to modulate the electrical activity of the targeted nerve fiber population. Therapeutic efficacy is directly influenced by electrode design, placement, and stimulus parameters. Computational modeling approaches have proven to be an effective way to select the appropriate stimulation parameters. Such an approach is, however, poorly accessible to inexperienced users as it typically requires the use of multiple commercial software and/or development in different programming languages. Here, we describe a Python-based framework that aims to provide an open-source turnkey solution to any end user. The framework we developed is based on open-source packages that are fully encapsulated, thus transparent to the end-user. The framework is also being developed to enable simulation of granular complexity, from rapid first-order simulation to the evaluation of complex stimulation scenarios requiring a deeper understanding of the ins and outs of the framework.

## Introduction

The network of peripheral nerves presents an extraordinary potential for modulating and monitoring the functioning of internal organs or the brain. Recent developments in electroceuticals showed promising results, highly targeted treatment, and fewer side effects when compared to more conventional drug-based therapies [1]. Electrical stimulation works by applying an electrical current via dedicated electrodes to a targeted neural tissue in the nervous system to generate or modulate the desired neural activity [2]. Currently, electrical stimulation is used in various applications in the Central Nervous System (CNS), such as Deep Brain Stimulation for Parkinson’s Disease [3] or for epileptic seizure reduction [4], and is also investigated in Peripheral Nervous System (PNS) for applications such as motor rehabilitation [5] of restoration of some involuntary or visceral functions [6] for instance. More recently, non-conventional stimulation delivering a specific electrical kilohertz frequency continuous waveform to the neural target has been used to block unwanted neural activity for pain reduction [7].

Although the literature reports many encouraging novel bioelectronic therapies, many of them fail to reach a higher level of technology readiness and don’t succeed in translating to the market [8]. Indeed, the sizeable morphological variety across species and within individuals of the same species greatly impacts the electrical stimulation therapy outcomes [9]. Also, the electrode design and its spatial location and orientation, as well as the choice of the electrical stimulus will further increase the lack of predictability of the therapy. Computational modeling has proven to be a valuable tool to tackle those issues but also for accelerating the design of the therapy while containing the financial and ethical cost of the *in-vivo* experiments [10–13]. In addition, *in-silico* models provide access to individual fiber responses and insight into the internal states of the neurons that constitute the simulated neural target. It helps to better understand the complex behavior of neural fibers and enables the design of specific therapies [14–16]. The process for simulating the PNS response to electrical stimuli is now rather consensual and is based on a 2-step hybrid model: [17, 18]:

1. A realistic 3D model is created. It includes the anatomical and physiological features as well as an accurate representation of the electrode design and location in or around the neural tissue. A Finite Element Modeling (FEM) solver is commonly used to compute the spatial extracellular electric potential distribution in the tissue induced by the electrical stimulation. This step usually relies on the quasi-static approximation [19].
2. The extracellular potential is then applied to unidimensional compartmental models that simulate the response of axon fibers to the electrical stimuli. In the case of PNS fibers, the ephaptic coupling between fibers is usually neglected or taken into account for very specific studies [20]. Hodgkin-Huxley-like (i.e. nonlinear conductance-based) models are commonly used, and distinct models are developed for specific trans-membrane protein and associated types of fiber (myelinated [21], unmyelinated [22], afferent/efferent [23], etc.).

Despite the benefits provided by hybrid *in-silico* studies, the approach remains complex to implement and is reserved for experts who are required to master several programming languages and/or software for the different steps of the simulation. Moreover, there is currently no framework that is universally used by the research community, reducing the possibility of experiment replications and the reuse of data [24]. In some cases, the published framework uses non-open and/or licensed software thus restricting their potential adoption by the community. Multi-software and/or multi-programming language approach also makes the deployment of hybrid modeling frameworks complex on the super-computer cluster or massively parallel machines.

In this paper, we propose and describe a fully open PNS simulation framework. This framework has been designed as a layer of abstraction of physical, mathematical, and computational techniques required for realistic prediction of bioelectronic phenomena prediction and study. It can be used with a few lines of code (in Python language) and used on various computing supports, from conventional machines to large supercomputers without impacting the simulation description. This framework has also been designed to be linked to optimization algorithms and experimental setups, to ease the translation between novel stimulation protocol development and experimental campaigns. The methods section start with a literature review of the existing framework, and then describes the architecture of the our framework, the algorithms, the physics, and the hypotheses behind the simulation with references to the existing literature. In the results section, we validate the output of the framework with existing results and show its potential applications. Finally, we discuss the advantages of our approach compared to the related literature and future development directions.

## Methods

### Overview of existing frameworks

An extensive description of the processes for obtaining and simulating hybrid models was first introduced by Raspopovic *et al*. and then refined by Romeni *et al*. [17, 18]. The authors provided a systematic methodology to build and exploit realistic hybrid models for designing PNS interfaces. The workflow was demonstrated using COMSOL Multiphysics (COMSOL AB, Stockholm, Sweden) and MATLAB (The MathWorks Inc, Massachusetts, United States) to build the geometry and solve the FEM. The neural dynamic was computed using NEURON [25]. Some algorithms and examples are made available on a public repository. However, no out-of-the-box framework is provided making the deployment of workflow challenging. Recently, however, a couple of frameworks based on the same or a similar methodology have been developed and introduced to the community to simplify the deployment of hybrid modeling. Those frameworks are introduced in this section.

#### PyPNS

PyPNS [26] is an open-source python-based framework aiming at modeling electrical stimulations and neural recordings of a nerve of the PNS. The MRG model [21] is used for simulating myelinated fibers, and the Sundt model [27] for unmyelinated ones. Neural dynamics are solved using the NEURON simulation software via the Python API [28]. Fiber models are stimulated via an extracellular electrical field that is precomputed using an external FEM solver. This framework separates time and space via the quasi-static approximation, limiting the need for multiple runs of the external FEM simulation. The PyPNS framework also aims at improving the simulation by taking into account the tortuosity of axons in the PNS nerve.

#### ASCENT

ASCENT [29] provides a computational modeling pipeline for the simulation of the PNS that very closely follows the methodology suggested in [18]. The pipeline uses Python and Java languages, COMSOL Multiphysics for the FEM problem, and NEURON for solving neural dynamics. Simulation parameters are defined in JSON configuration files and the simulation is a 2-step process: i) Python, Java, and COMSOL Multiphysics are used to evaluate the potentials along neural fibers. ii) Python and NEURON are used to compute the resulting neural activity. The second step can easily be run on a computer cluster. A standardized pipeline is developed to describe and process nerve morphology using the Python language. Histology-based or *ex-novo* nerves can be easily created. Arbitrary waveforms can be generated enabling the simulation of neural activation and neural block. The pipeline provides tools for analysis and visualization such as heatmaps of thresholds or video generators for plotting state variables variation over time and space. The pipeline is freely available on a repository, and extensive examples and documentation are provided but requires a commercial licence.

#### TxBDC Cuff Lab

The TxBDC Cuff Lab [30] is a Python web server with an online front-end graphical user interface that can run simple parametrized simulations to estimate electrical stimulation’s recruitment curves. The geometry and electrical properties of the nerves can be adjusted. Electrodes can be either cuff-like or intraneural electrodes and can be also parameterized. Stimulation waveform is limited to either a monophasic or a biphasic pulse. The FEM model is meshed with Gmsh [31] and solved under the quasi-static assumption using the FEniCS package [32]. Neural dynamics are computed with passive axon models. The model is based on 3-D fitted curves obtained from a pre-computed active axon model modeled with NEURON. A total of 20 fitted curves are pre-computed and used to estimate neural dynamics. Backed-end Python sources are freely available on a dedicated repository.

#### ViNERS

ViNERS [33] is an open-source multi-domain MATLAB-based toolbox that models visceral nerves. Similarly to PyPNS, ViNERS can simulate electrical stimulation and neural activity recordings. FEM is done internally using Gmsh for meshing the geometry and is solved with EIDORS [34]. Both stimulation and recordings use FEM providing realistic outputs. Neural dynamic is solved via NEURON. Nerve design and electrode layout can be done via scripting or a JSON configuration file. The JSON file can be generated with a dedicated graphical user interface (GUI). Nerve anatomy can be imported from a traced nerve section from a histology sample or *ex-novo* anatomy can be created. Un-myelinated/myelinated efferent and afferent axons are populated from measured fiber distribution. MRG and Gaines [23] models are used for myelinated fibers and the Sundt model for unmyelinated fiber. Axon structural parameters are extrapolated from their original values to model small axon diameters present in visceral fascicles correctly. ViNERS also provides tools to analyze simulation output such as spiking threshold detection or single-fiber action potential detection for recordings. ViNERS is provided as a MATLAB toolbox that can be freely downloaded. Source code and examples are also provided. ViNERS is, to the best of our knowledge, the most complete open-source framework available, as all the required steps can be done within the toolbox.

#### Sim4Life

Sim4Life (Zurich Med Tech, Zurich, Switzerland) is a multi-physics and multi-domain commercial simulation platform. It includes FEM and neural simulations, a 3D modeling environment as well data analysis and visualization tools. Human and animal-validated 3D models are provided and the platform is optimized for computational performances. Each step of the simulation can be parameterized via a GUI and no programming language knowledge is required. The software is provided with examples, documentation, and access to customer service.

The development of dedicated frameworks for hybrid modeling is a great step forward towards simple, efficient, and easy-to-replicate *in-silico* electrical stimulation of the PNS. However, the presented frameworks suffer from several limitations. For example, Sim4Life is a licensed software thus limiting its accessibility. Also, it is a multi-physics simulator not specialized in the simulation of the PNS. It comes at the cost of a relative complexity and some training might be required. Its close-source approach limits the possibility of customization and extension, as well as restraints the comprehension of models utilized. ASCENT and ViNERS are open-source frameworks but require licensed software to run: ViNERS is MATLAB-based, and ASCEND requires COMSOL Multiphysics to run. PyPNS does not include the FEM solver and the FEM was pre-computed using COMSOL Multiphysics. The TxBDC Cuff Lab approach is fully open-source and easy to use but is intrinsically limited to simplified nerve and electrode geometries as well as stimulation waveforms.

### Our approach: NRV Overview

The NeuRon Virtualizer framework (NRV) is a fully open-source multi-scale and multi-domain Python-based framework developed for the hybrid modeling of electrical stimulation in the PNS. NRV shares the same working principle as the frameworks presented in the previous section: a FEM model is solved under the quasi-static assumption to compute the extracellular potential generated by a source and is applied to a neural target. However, NRV improves the previous frameworks with its capability of performing all the required steps for hybrid modeling within the framework. Realistic geometries for the electrodes, nerves, and fascicles are constructed and meshed using Gmsh [31], and FEM problems are solved with the FEniCS software [32]. This workflow is fully open-source and requires no software purchase from the end user. A bridge between COMSOL Multiphysics and NRV via COMSOL Server (requiring a commercial license) is also implemented in NRV providing multiple options to the users for creating the 3-D model. Tools for generating realistic axon population and placement are provided and commonly used in the literature axon models are implemented.

Extracellular potential computed with the FEM model is interpolated and used as input for the 1-D axon models. An intracellular voltage or current clamp can also be used to stimulate one or multiple axons of an NRV model. Axons are then simulated using the NEURON framework with the NEURON to Python bridge. All computation inputs and outputs are stored in dictionary objects to enable context saving. Post-processing tools are also provided to automatically detect spikes, filter the data, etc. Extracellular recorders are also implemented and can be added to the model to simulate electrically-evoked compound action potentials (eCAPs). Large-scale nerve simulations are embarrassingly parallel problems and can greatly reduce computation time. NRV provides multiprocessing capabilities by implementing a message-passing interface (MPI) for Python [35]. Parallel computing is performed independently from the simulation description and the end-user only needs to provide the maximum number of usable cores on the machine; NRV automatically handles job distribution and synchronicity between the processes. Calls to Gmsh, FenicsX, COMSOL Server, NEURON, or other third-party libraries used are fully encapsulated in the NRV framework making their use completely transparent to the user.

NRV aims at being accessible for users with only basic Python knowledge as well as easily readable from high-level simulation perspectives. NRV also enables multi-scale simulations: single axonal fibers to whole nerve simulations can be performed with NRV and require only a couple of lines of code. The framework is pip installable making the framework effortlessly deployable on a computer cluster or supercomputer.

NRV’s internal architecture is depicted in Fig 1. It is subdivided into four main sections:

**Fig 1.**
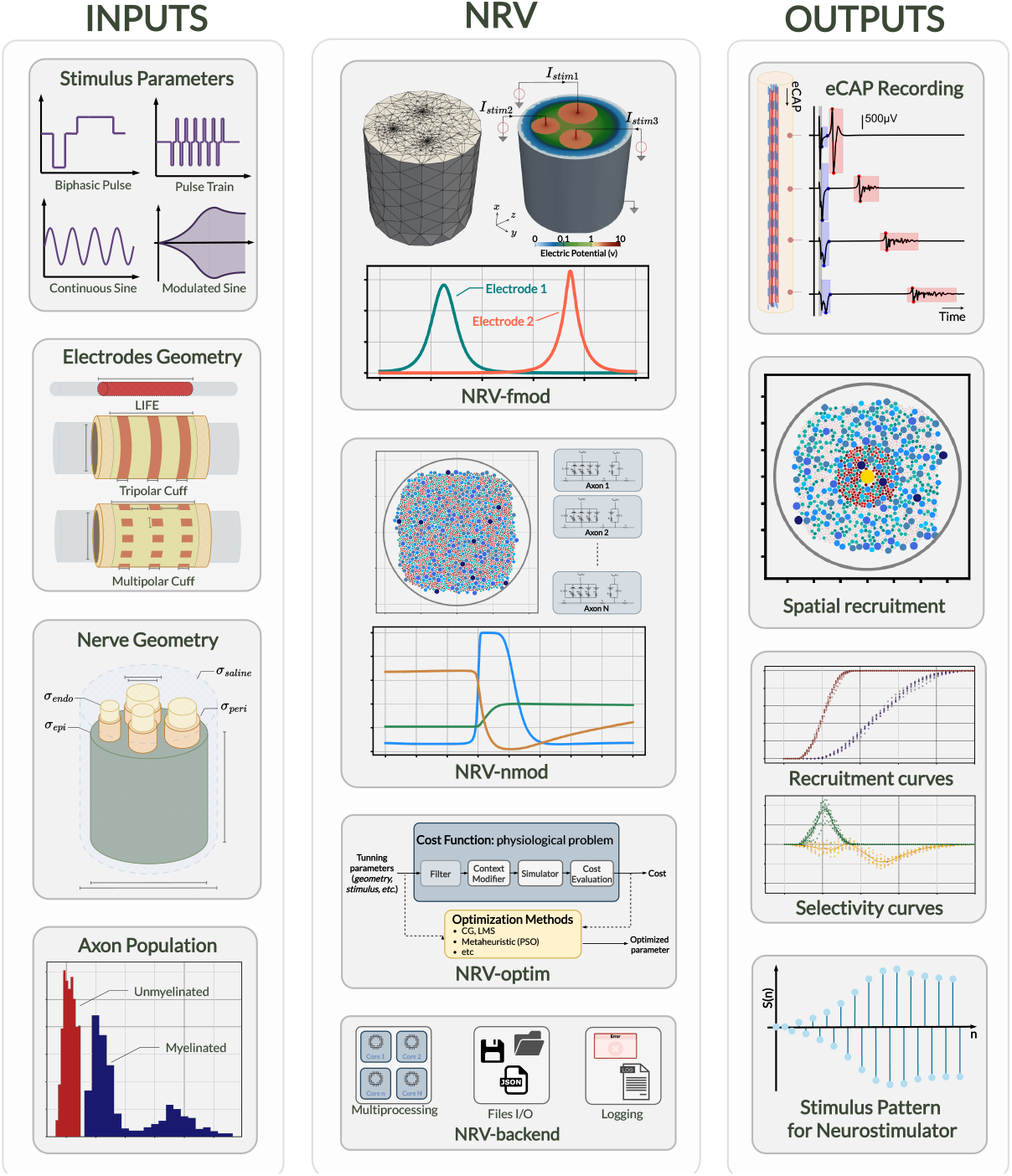
Schematic overview of NRV. The figure shows the framework inputs (stimulus parameters, electrode and nerve geometry, and axon populations), the framework mains sections (*fmod, nmod, optim*, and *backend*) described in the **Method** section of this paper, and some possible result outputs of the NRV framework.

- <u>fmod:</u> handles 3-D extracellular model generation and computation.
- <u>nmod:</u> manages 1-D axon membrane potential model description and computation.
- <u>optim:</u> enables automated optimization of stimulation contexts, either by controlling geometrical parameters or the stimulation waveshape for instance.
- <u>backend:</u> manages all related software engineering aspects behind NRV, such as machine capacity and performance, parallel processing, or file input and output. This section is not related to the neuroscientific computational aspect and is not described further below.

### Fmod-section: computation of 3D extracellular potentials

NRV provides classes, tools, and templates to create 3D models of the nerve and electrodes in the Fmod-section of the framework. Fig 2 provides a synthetic overview of the extracellular simulation problem and a simplified UML-class diagram of the software implementation, including class references accessible to the end user. In this paragraph, we provide details of the implementation of the physics and the corresponding computational mechanisms.

**Fig 2.**
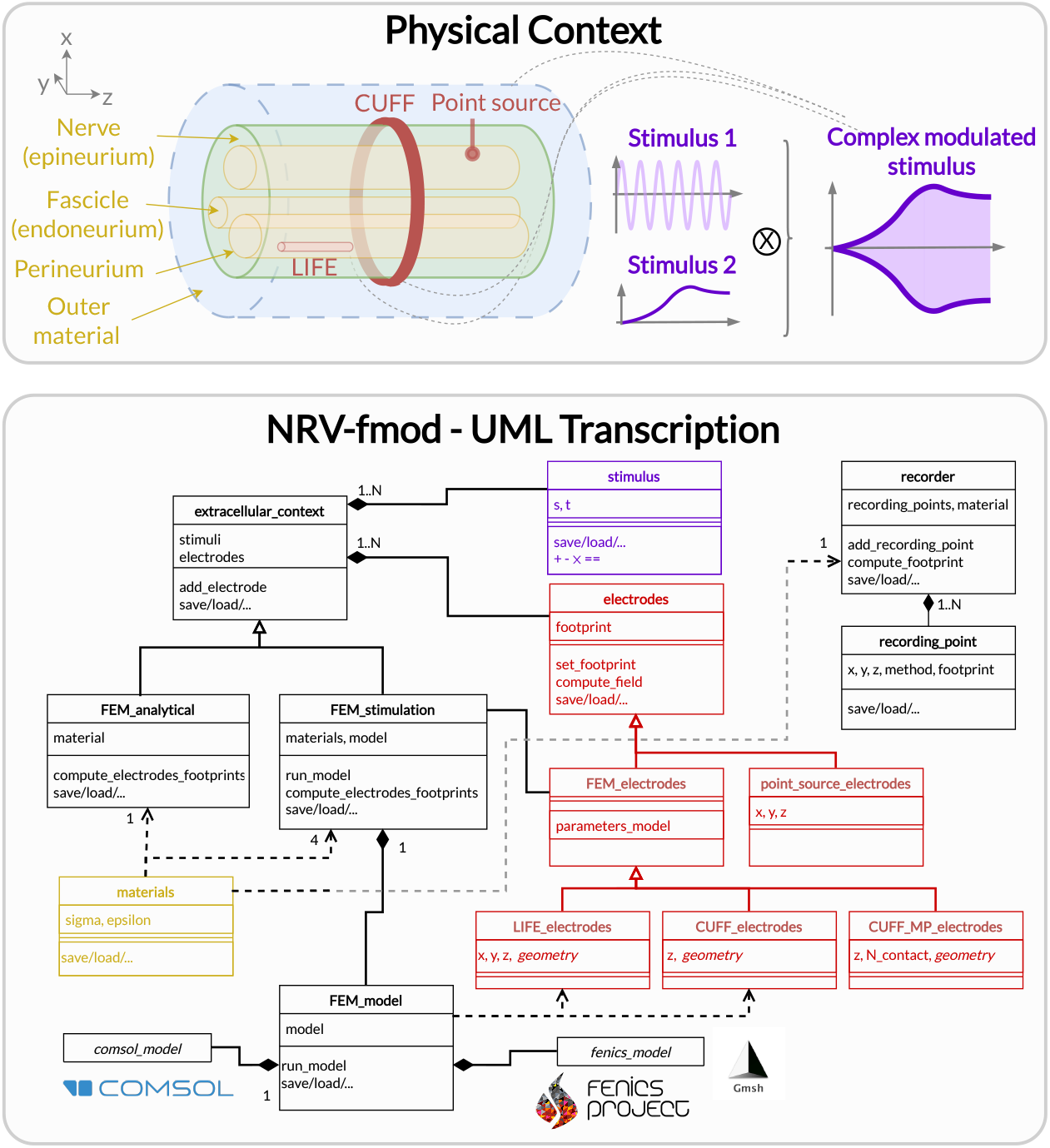
Fmod section of the NRV framework. Top: Schematic conceptualization of a PNS extracellular simulation context. The context is created around a nerve, its fascicles, and corresponding materials. Electrodes can be added as electrical interfaces, each electrode contact being associated with a simple or complex stimulus (or stimulation waveform). Bottom: NRV’s transcription of the physical with dedicated classes that can be combined and are used by the framework to evaluate the extracellular electrical fields.

The estimation of the extracellular electric potential resulting from the electrical stimulation is handled by the `extracellular_context`-class. The `analytical_stimulation`-class and the `FEM_stimulation`-class are derived from the parent `extracellular_context`-class as illustrated in Fig 2 and detailed in the next two subsections. The extracellular potential generated by neural activity is handled by the `recorder`-class and described later on.

#### Electrical stimulation potential: computation mechanism

NRV relies on the assumption of linear impedance material, so that the contribution of each electrical source, i.e. stimulating electrode, can be linearly combined. In addition, the quasi-static approximation leads to a time-decoupled solution of the electrical stimulation potential. As so, for a point of coordinate **r** in space (**r** *∈* ℝ^3^) at a time *t*, the extracellular electrical potential is computed as:

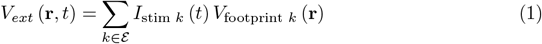

where *ℰ* is the set of electrodes in the simulation, *I*_stim *k*_ is the stimulation current at the electrode *k*, and *V*_footprint *k*_ (**r**) is a function of ℝ^3^ → ℝ computed once for each electrode before any simulation and which corresponds to the 3D extracellular electrical potential generated by the electrode for a unitary current contribution. This electrode contribution function can be computed using two different methods, explained below, and the result is referred to in this article as the electrode footprint.

The stimulation current *I*_stim,*k*_ is described by a dedicated `stimulus`-class that defines asynchronous stimulus current and time values applied to an electrode *k*. Arithmetic operations between `Stimulus`-objects are defined for the class, thus enabling fully customized arbitrary stimulating waveforms. This approach facilitates the design of complex waveforms such as modulated stimuli [36–38]. NRV also includes dedicated methods for fast generation of monophasic and biphasic pulses, sine-wave, square-wave, and linear ramps stimulus.

When describing the stimulating electrode by instantiating an `extracellular_context`-object, the end user can choose between two different methods to compute the extracellular potential: an analytical approach or a FEM approach. This choice affects not only the computational requirements but also the degree of realism of the simulation. As described in the class diagram of Fig 2, the end user can add a combination of electrodes (`electrode`-class) and stimuli (`stimulus`-class) no matter the extracellular potential computation approach selected. Each `Electrode`-object in NRV has a unique ID and multiple electrodes can be added to the simulation model. If the extracellular potential is solved analytically, only Point-Source Approximation (PSA) electrodes can be implemented. Using the FEM approach, classes to simulate cuff-like electrodes and longitudinal intrafasicular electrodes (LIFEs) [39] have been implemented. FEM electrodes can be fully parameterized (active-site length, number of contacts, location, etc.). Custom classes for more complex electrode designs can be further implemented by inheritance of the `FEM_electrodes`-class. All footprint computations are performed by the `electrode`-mother class automatically when the `extracellular_context`-object is associated with axons (mechanism described in the next section, see Fig 4).

**Fig 3.**
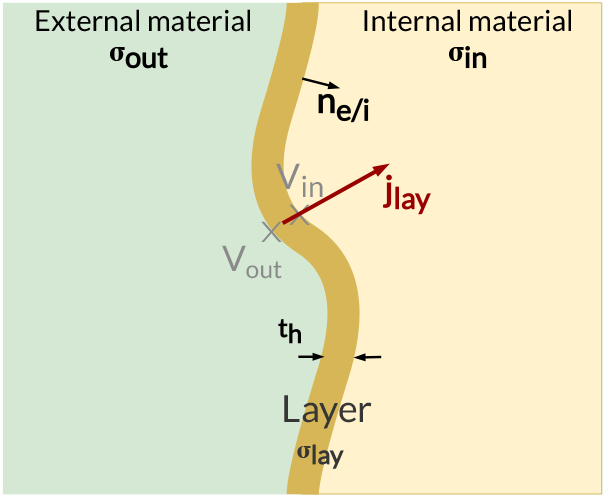
Thin-layer approximation in NRV. Thin-layer of thickness *t*_*h*_ and conductivity *σ*_*lay*_, bounding two materials, internal and external, of conductivity *σ*_*in*_ and *σ*_*ext*_ respectively.

**Fig 4.**
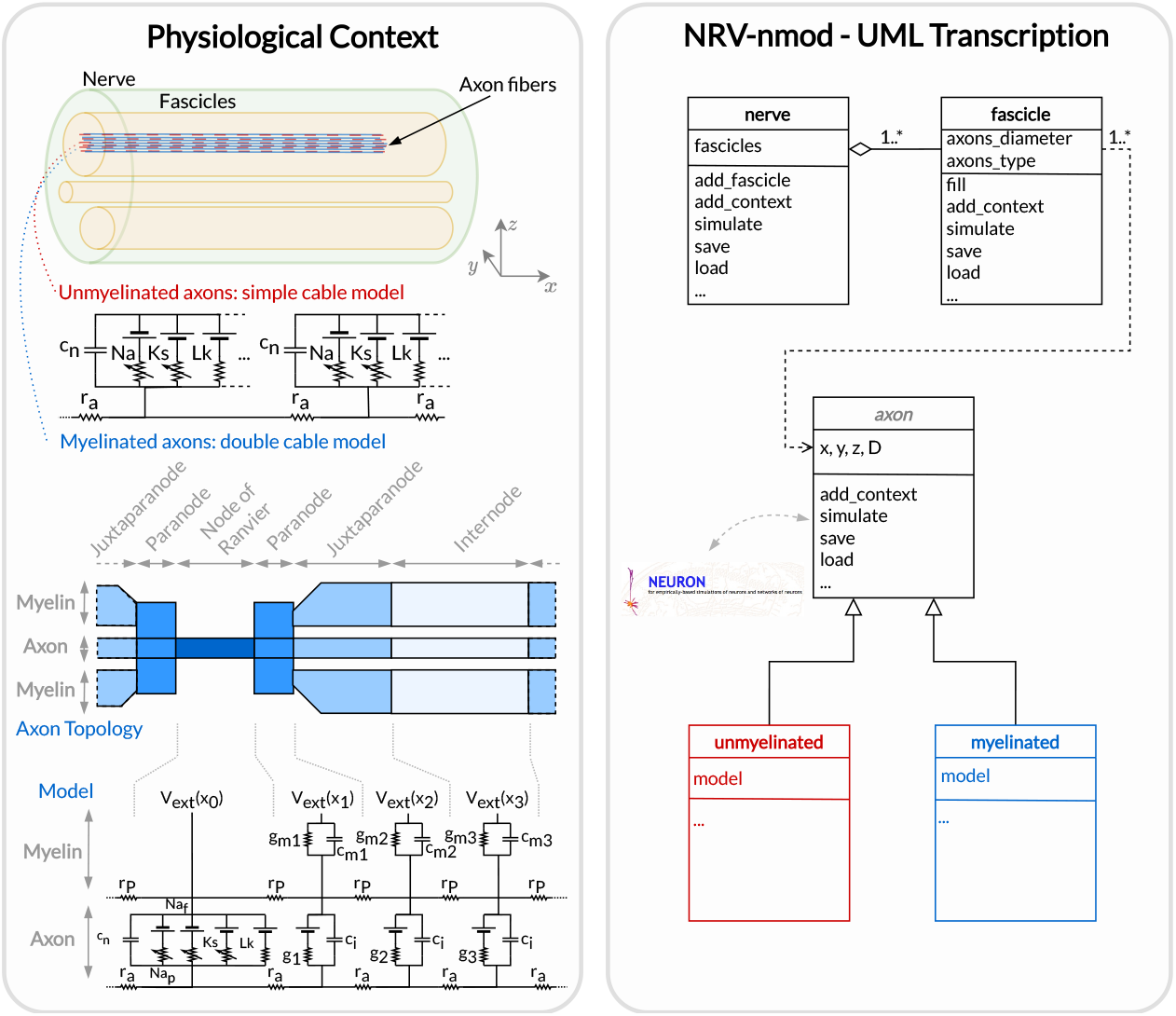
Node section of the NRV framework. Left: Overall view of the neuronal context of a generic PNS simulation. A nerve is an entity composed of one or multiple fascicles that do not overlap, each containing myelinated and/or unmyelinated axons. Right: NRV’s transcription of the neuronal context with dedicated classes that can be combined and used by the framework to evaluate the resulting neural activity of each axon.

Electrical conductivities (isotropic or anisotropic) of the tissues constituting the NRV nerve are defined using `Material`-class. The framework includes pre-defined materials for the epineurium, endoneurium, and perineum conductivities with values commonly found in the literature [40]. Custom conductivity values can also be user-specified for each material used in the simulation.

#### Analytical evaluation of the extracellular potential

The `analytical_stimulation`-class solves the extracellular potential analytically using the PSA for the electrode and the nerve is modeled as an infinite homogeneous medium [41]. This method is only suitable for geometry-less simulation: axons are considered as being surrounded by a unique homogeneous material. In this case, the footprint function is computed as:

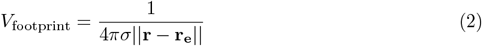

where || *·* || denote the euclidean norm, **r**_**e**_ is the (*x*_*e*_, *y*_*e*_, *z*_*e*_) position of the PSA electrode and *σ* is the isotropic conductivity of the material. The conductivity of the endoneurium is anisotropic as it is higher in the radial orientation of axons [40]. For anisotropic material, the conductivity becomes a diagonal matrix:

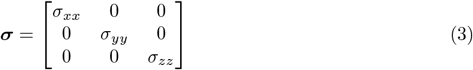

The expression of the footprint function becomes [42]:

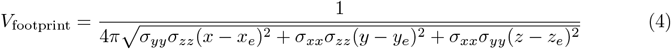

The analytical approach provides a simple and quick estimation of the extracellular potential, enabling fast computation on limited-resource machines. However, it restricts the geometry to only one material and the electrode shape is not taken into account, limiting the viability of this approach to model complex experimental or therapeutic setups [43].

#### FEM evaluation of the extracellular potential

The extracellular potential evaluation in a realistic nerve and electrode model using the FEM approach is handled by the `FEM_stimulation`-class. A nerve in NRV is modeled as a perfect cylinder and is defined by its diameter, its length, and the number of fascicles inside. The position and diameter of each fascicle on the NRV nerve can be explicitly specified. Fascicles of the NRV model are modeled as bulk volumes of endoneurium surrounded by a thin layer of perineurium tissue [44]. The remaining tissue of the nerve is modeled as a homogeneous epineurium. The nerve is plunged into a cylindrical material, which is by default modeled as a saline solution.

The NRV framework offers the possibility of using either COMSOL Multiphysics or FEniCS to solve the FEM problem. For the first one, mesh and FEM problems are defined in `mph` files which can be parameterized in the `FEM_stimulation`-class to match the extracellular properties, and all physic equations are integrated into the `Electric Currents` COMSOL library. For the FEniCS solver, NRV handles the mesh generation using Gmsh, the bridge with the solver, and the finite element problem with FEniCS algorithms. Physic equations solved are defined within the NRV framework. COMSOL Multiphysics is commonly used for the simulation of neural electrical stimulation investigation, but it requires a commercial license to perform computation, and all future developments are bound to the physics and features available in the software. We included the possibility of using it as a comparison to existing results but the use of FEniCS and Gmsh enables fully open-science and the possibility to enhance simulation possibilities and performances.

For both FEM solvers, the extracellular electrical potential *V*_footprint_ in the simulation space Ω, is obtained from the current conservation equation under the quasi-static approximation and the structural equation in a conductive material of conductivity *σ*:

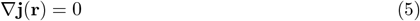

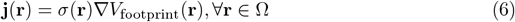

where **j** is the current density. The electrical ground and the current injected by an electrode are set by respectively Dirichlet and Neumann boundary conditions:

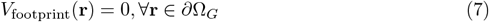

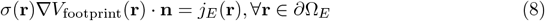

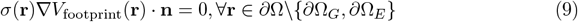

where the ground surface *∂*Ω_*G*_ can be specified by the end user in one or multiple external surfaces of the extracellular space. At the electrode interface, **n** is the normal vector to the surface *∂*Ω_*E*_ and the normal current density injected *j*_*E*_ considered homogeneously distributed is computed by:

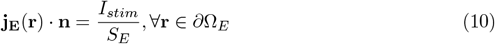

where *I*_*stim*_ is the stimulation current and *S*_*E*_ the electrode contact surface area.

To reduce the number of elements in the mesh associated with smaller material dimensions, the fascicular perineurium volumes are defined using the thin-layer approximation (see Fig 3) [44, 77]. The current flow is assumed to be continuous through the layer, while a discontinuity is induced in the potentials:

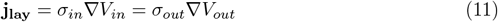

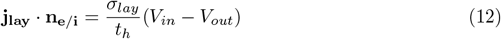

#### Simulation of eCAP recordings: computation mechanism

Using the linear material impedance hypothesis, the extracellular electrical potential can be considered as the sum of the contribution of stimulation electrodes and electrical potential generated by axons’ neural activity. As so, the two contributions can be computed separately. With the hypothesis of no ephaptic coupling, the contribution of each axon’s action potential is considered negligible compared to the potential due to the stimulating electrodes. Therefore the axon activities are evaluated independently from each other.

In NRV, we chose to compute eCAPs using first approximation hypotheses: eCAPs are computed analytically only, using a point- or line-source approximations (PSA or LSA) [45] of the contribution of each axon in the simulation. The geometry is thus only based on one material (by default endoneurium). This strategy ensures computational efficiency while still providing sufficiently quantitative results about axon synchronization and eCAP propagation for comparison with experimental observations.

The eCAP recording is performed automatically for the user when instantiating a `recorder`-object (see Fig 2), linking one material with multiple `recording-points`-objects. These last objects are simply positions in space where the extracellular is recorded for the duration of the simulation. Using again space and time decoupling, the eCAP electrical potential at a position **r** at a time *t* is computed as:

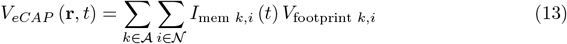

where 𝒜 is the set of axons in the simulation, *𝒩* is the set of computational nodes in the axon implementation (see Nmod section below), *I*_mem *k,i*_ the membrane current computed in the nmod section(see below) and *V*_footprint *k,i*_ is a scalar. From a numerical perspective, equation 13 is equivalent to a sum of dot products between two vectors: the membrane current computed in the nmod section of NRV (see below) and a recorder footprint. The footprint is computed only once for each axon in the nerve geometry before any simulation.

The footprint for one position **r**_**k**,**i**_ = (*x*_*k,i*_, *y*_*k,i*_, *z*_*k,i*_) *∈* ℝ^3^ in space corresponding to the node *i* of the axon *k* for a `recording-points`-object at the position

*r*_*rec*_ = (*x*_*rec*_, *y*_*rec*_, *z*_*rec*_) *∈* ℝ^3^ is computed either with PSA:

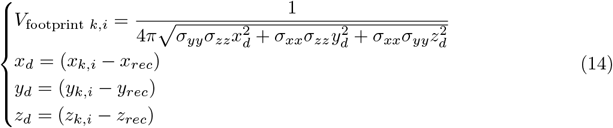

for anisotropic or isotropic materials (*σ* = *σ*_*xx*_ = *σ*_*yy*_ = *σ*_*zz*_), of with LSA for isotropic materials only [45]

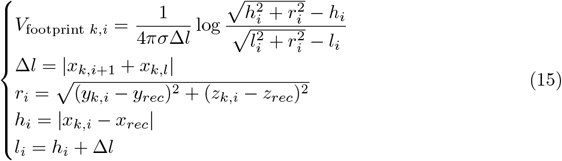

In both cases, eCAP computation is performed after the computation of neural activity, which is explained in the next section.

### Nmod section: generating and simulating axons

The description of a neuronal context in NRV and the computation of the trans-membrane potential are described in a hierarchical manner in Fig 4. At the bottom of the hierarchy, axons are individual computational problems for which the NRV framework computes the membrane voltage response to extracellular electrical stimulation. The conventional hypothesis is that each axon is independent of others, i.e. there is no ephaptic coupling between fibers, so all axon computations can be performed separately. From a computational point of view, this hypothesis transforms neural computation into an embarrassingly parallel problem, allowing massively parallel computations. In this section, details of the models are given using a bottom-up approach: axon models are described first, followed by fascicle entities, and finally nerves.

#### Axon models

Axonal fibers in NRV are defined with the `axon`-class. This class is an abstract Python class and cannot be called directly by the user. It however handles all generic definitions and the simulation mechanism. Axons are defined along the *x − axis* of the nerve model. Axon coordinates and axon length are specified at the creation of an `axon`-object. End-user accessible `Myelinated`-class and `unmyelinated`-class define myelinated and unmyelinated fiber objects respectively and inherit from the `axon`-class.

Computational models can be specified for both the myelinated and unmyelinated fibers. Currently, NRV supports the MRG [21] and Gaines [23] models for myelinated fibers. It also supports the original Hodgkin-Huxley model [46], the Rattay-Aberham model [47], the Sundt model [27], the Tigerholm model [22], the Schild model [48] and its updated version [49] for unmyelinated fibers.

MRG and Gaines model’s electrical properties are available on ModelDB [50] under accession numbers 3810 and 243841 respectively. Interpolation functions used in [23] to estimate the relationship between fiber diameter and node-of-Ranvier, paranode, juxtaparanodes, internode length, and axon diameter generate negative values when used with small fiber diameter. In NRV, morphological values in [21] and from [51] are interpolated with cubic or quadratic functions. The juxtaparanode length is fitted with a 5^*th*^ order polynomial function between 1*μm* and 16*μm* and with a linear interpolation outside this range. Parameters of the unmyelinated models are taken from [52] and are available on ModelDB under accession number 266498.

The extracellular stimulations are connected to the `axon`-object with the `attach_extracellular_stimulation`-method, which links the `extracellular_context`-object to the axon. Voltage and current Patch-clamps can also be inserted into the axon model with the `insert_V_Clamp`-method and `insert_I_Clamp`-method. The `simulate`-method of the `axon`-class solves the axon model using the NEURON framework. NRV uses the NEURON-to-Python bridge [28] and is fully transparent to the user. The `simulate`-method returns a dictionary containing the fiber information and the simulation results.

#### Fascicle construction and simulation

The `fascicle`-class of NRV defines an individual population of fibers. The `fascicle`-object specifies the number of axons in the population, the fiber type (unmyelinated or myelinated), the diameter, the computational model used, and the spatial location of each axonal fiber that populates the simulated fascicle.

An axon population can be pre-defined and loaded into the `fascicle`-object. Third-party software such as AxonSeg [53] or AxonDeepSeg [54] can be used for generating axon populations from a histology section that are then loaded into the `fascicle`-object. Alternatively, the NRV framework provides tools to generate a realistic *ex-novo* population of axons. For example, the `create_axon_population`-function creates a population with a specified number of axons, a proportion of myelinated/unmyelinated fibers, and statistics for unmyelinated and myelinated fibers’ diameter repartition. Statistics taken from [55–57] have been interpolated and predefined as population-generating functions. User-defined statistics can also be specified. Alternatively, the `fill_area_with_axons`-function fills a user-specified area with axons according to the desired fiber volume fraction, fiber type, and diameter repartition statistics. An axon-packing algorithm is also included to place fibers within the fascicle boundaries. The packing algorithm is inspired by [58]. The generation of a realistic population and the packing principle are illustrated in Fig 5.

**Fig 5.**
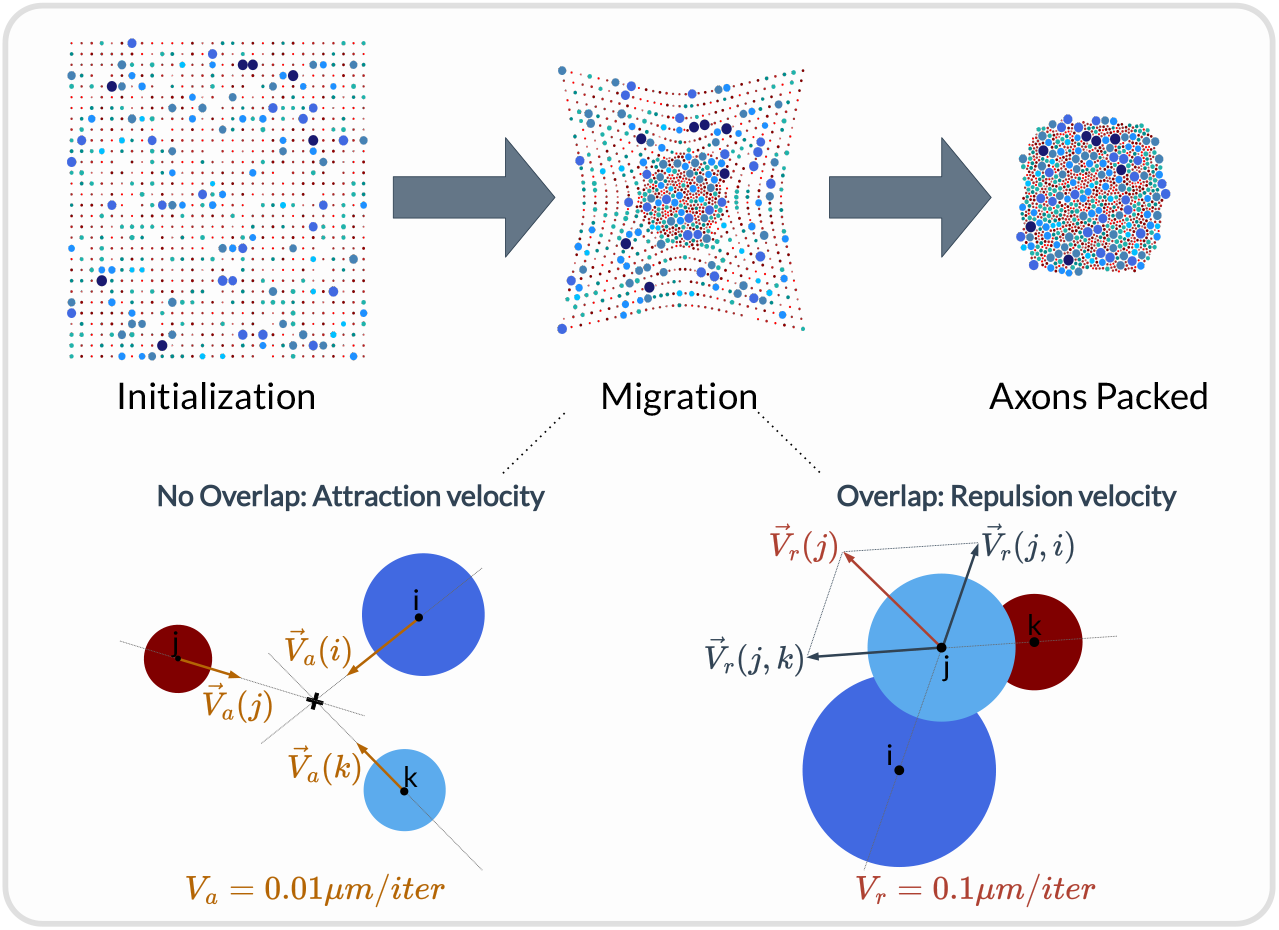
Overview of the axon-packing algorithm inspired from [58]. The packing procedure was demonstrated with 1000 fibers. Myelinated axons are in blue and unmyelinated ones are in red. Fibers are randomly placed on a grid at initialization and migrate progressively toward the center. Each axon’s new position is calculated at each iteration according to an attraction velocity or a repulsion velocity if axons are overlapping.

The `fascicle`-class can perform logical operations (remove geometrically overlapping populations, diameter filtering) and geometrical operations (translations and rotations) on the axon population. Node-of-Ranvier of the myelinated fiber can be also aligned or randomly positioned in the fascicle. An `extracellular_context`-object is added to the `fascicle`-object using the `attach_extracellular_stimulation`-method. Intracellular stimulations can also be attached to the entire axon population or to a specified subset of fibers. The `simulate`-method creates an `axon`-object for each fiber of the fascicle, propagates the intracellular and extracellular stimulations and recorders, and simulates each of them. Parallelization of `axon`-object simulation is automatically handled by the framework and fully transparent to the user. The simulation output of each axon is saved inside a pre-defined folder.

#### Simulation top level: the Nerve-object

To achieve realistic PNS simulation, a top-level `nerve`-class is implemented. Nerves typically consist of one or more fascicles. Therefore, class aggregation allows logical and geometric operations between contained fascicles. In addition, `extracellular_context`-objects and intracellular stimulations can be added globally. The simulation is handled automatically by the framework and thus, as for `fascicle`-objects, is fully transparent to the end-user.

The geometrical parameters of the `nerve`-object are used for automatic generation of the geometry of the physical context description (see Fig 2), and all extracellular context simulations are triggered automatically and fully transparent to the user.

### Optimize

Simulation frameworks such as NRV are often used in an open-loop manner: the end user defines a context corresponding to an electro-physiological setup (nerve physiology, electrode geometry, stimulation sites, and waveforms, recording) and explores quantities of interest by screening some parameters over a certain range. This process enables improvements in stimulation performance at the cost of a long and fastidious process. More recently, the idea of using optimization algorithms in a closed-loop fashion with simulation models has been successfully applied to improve electrode design [59] or stimulation waveforms or protocol [36, 60, 61]. However, such methods require an important phase of formalization and software development. To ease and automatize such methods, the NRV framework includes an out-of-the-box solution for describing any optimization problem for PNS simulation. Fig 6 describes the adopted generic formalism: an optimization problem (defined in a `Problem`-class) couples a `Cost_Function`-object in charge of evaluating the impact of some parameters (included in a vector) on a simulation, and an optimization method or algorithm embedded in the `Optimizer`-object. Optimization methods rely on third-party optimization libraries: SciPy optimize [62] for continuous problems, Pyswarms [63] as Particle Swarms Optimization metaheuristic for high-dimensional or discontinuous problems.

**Fig 6.**
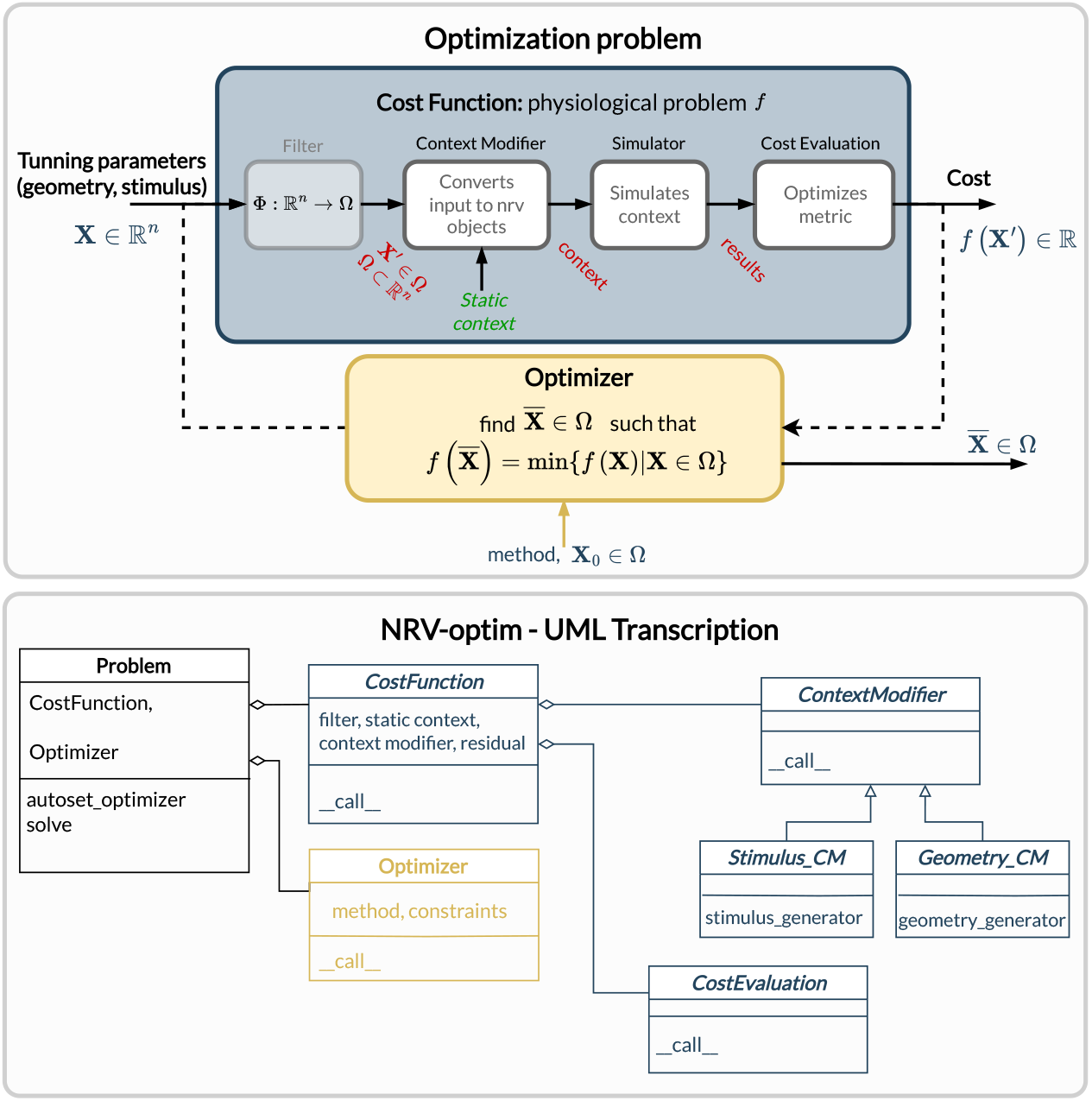
Overview of optimization problem formalism and implementation in NRV. A problem is described by combining a cost function and an optimizer or optimization method. The cost function transforms a vector to a cost by modifying any static simulation (of an axon, fascicle, or nerve) and computing a user-defined cost from the simulation result. Corresponding classes have been developed to ease the formulation of problems.

The novelty provided by our framework is the ease of gathering elements required to evaluate the impact of a specific set of parameters associated with a simulation and a final cost evaluation. The `Cost_Function`-class is constructed around (referring to Fig 6):

- A filter: which is an optional Python `callable`-object, for vector formatting or space restriction. In most cases, this function is set to identity and will be taken as such if not defined by the user.
- A static context: it is the starting point of the simulation to be optimized. It can be one of the NRV-nmod objects (axon, fascicle, or nerve) to which all objects describing stimulation, recording and more generally the physical context are attached. These NRV-nmod objects have in common a `simulate`-method.
- A `ContextModifier`-object: it generates an updated local context from the static context and input vector. The `ContextModifier`-object is an abstract class, and two classes of problems are currently predefined: problems for stimulus waveform optimization or for geometry (mainly electrodes) optimization. However, there is no restriction to define any exotic optimization scenario by inheriting from the parent `ContextModifier`-class.
- A `CostEvaluation`-object: uses the simulation results to evaluate a user-defined cost. Some examples of cost evaluation are included in the release framework. Nevertheless, the `CostEvaluation`-class is a generic Python `callable`-class so it can also be user-defined.

## Results

### FEM models cross-validation

While COMSOL Multiphysics is frequently used in hybrid modeling [17, 18, 26, 29], only the TxBDC Cuff Lab framework is based on the FEniCS solver [30]. The implementation of the FEM equations and the thin-layer approximation are first validated with a passive bi-domain 2-D model. Details and results of the validation are provided in the supplementary material (S1. Appendix). The remainder of this section evaluates the impact of the selected FEM solver (FEniCs or COMSOL Multiphysics) on the calculation of the electrode footprint and the estimation of the resulting axon’s activating thresholds. Python scripts for generating and plotting the data presented in this section are available in the supplementary material (S2. Files).

#### Electrode Footprint Computation

The extracellular voltage footprint generated by a LIFE and by a monopolar circular cuff-like electrode along an axonal fiber located at the center of a monofascicular nerve are compared with results obtained with COMSOL Multiphysics. LIFE active-site is 1000*μm* long and 25*μm* in diameter. Cuff-like electrode contact width and thickness are 500*μm* and 100*μm* respectively. The LIFE active site is placed 100*μm* away from the fiber inside the fascicle. The cuff electrode fully wraps the nerve and has a thin insulating layer on its outer surface to restrict current flow. The extracellular voltage footprint along the fiber, as well as its spatial second derivative, are shown in Fig 7(a) and (b) for the LIFE and the cuff-like electrode respectively. The relative difference between the two solvers’ estimation is also plotted.

**Fig 7.**
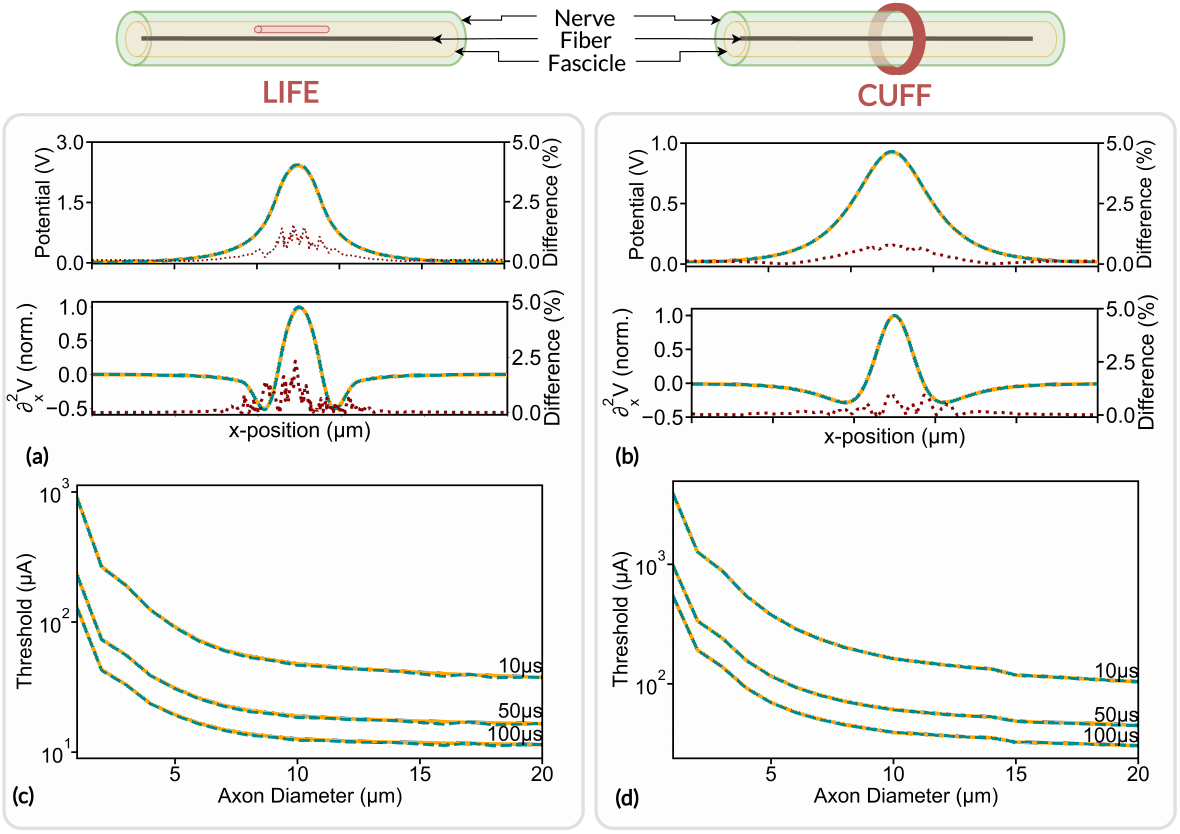
Electrode footprints and axon activation thresholds estimation. Footprints and activation thresholds are evaluated using the FEniCS solver (orange line) and COMSOL Multiphysics (dashed blue line) with a LIFE ((a) and (c)) and a cuff-like electrode ((b) and (d)). The relative difference in electrode footprint estimation is plotted in a red dotted line. The nerve and the fascicle are 1000*μm* and 800*μm* in diameter respectively. The fascicle is surrounded with a 5*μm* thick sheath of perineurium and the nerve is plunged in a 10*mm* large saline bath (not represented). The fascicle is made of anisotropic bulk endoneurium (*σ*_*x*_ = 0.57*S/m, σ*_*y*_ = 0.085*S/m*) and epineurium (*σ* = 0.085*S/m*). Perineurium and saline conductivity are set to *σ* = 0.002*/m* and *σ* = 2*S/m* respectively [40].

COMSOL Multiphysics and FEniCS estimated footprints are in very good agreement for both electrodes. The maximum electrode footprint difference is about 1.4% with LIFE and about 0.8% with a cuff-like electrode. The difference peaks at the maximum of the footprints, i.e. where the axonal fiber is the closest to the electrodes. Dissimilarities near the electrodes’ active site are to be expected as both FEM solvers are likely to handle boundary conditions slightly differently. The induced neural activity in axonal fiber is related to the spatial second derivative of the electrode footprint [47], which differs by a maximum of 2% with LIFE and a maximum of 1% with cuff electrode between COMSOL multiphysics and FEniCS estimations.

#### Activation Thresholds

This section evaluates the impact of the difference in electrode footprint estimation between FEniCS and COMSOL Multiphysics on the induced neural dynamics. Specifically, the influence on the activation threshold, i.e. the minimum stimulation current required to induce an action potential within an axonal fiber, is evaluated. The axonal fiber is modeled as a myelinated fiber using the MRG model [21] located at the center of the monofascicular nerve. The activation threshold is evaluated for an axon diameter ranging from 1*μm* to 20*μm*. The axon is stimulated with a monophasic pulse with a pulse duration of 10*μs*, 50*μs*, and 100*μs* delivered to the nerve with a LIFE and a cuff-like electrode. LIFE and cuff-like electrode footprints were computed with both COMSOL Multiphysics and FEniCS and the activation thresholds were estimated with the binary search algorithm implanted in the framework and with a search tolerance of 0.1%. Results are illustrated in Fig 7 (c) and (d) for the LIFE and cuff-like electrode respectively.

Thresholds estimated with FEniCS match very well those estimated with COMSOL Multiphysics, for both the LIFE and the cuff electrode. The maximum threshold estimation difference is about 1% the LIFE and about 0.4% with the cuff electrode. Mean differences are about 0.2% and 0.05% respectively which confirms the correct implementation of FEniCS in NRV. This also demonstrates that simulation results obtained with the two solvers can be safely compared and that the transition from COMSOL Multiphysics to FEniCS has a negligible effect on simulation results.

### Replication of previous results

#### Replication of *in-silico* studies

The NRV framework aims at facilitating the reproduction of *in-silico* studies. As an example, we replicated in this section a study on the high-frequency alternating current (HFAC) block mechanism modeled by Bhadra *et al*. [15]. The study demonstrated the phenomenon of HFAC nerve conduction block in a mammalian myelinated axon model and characterized the effect of HFAC frequency, electrode-to-axon distance, and fiber diameter on the HFAC block threshold. The axon fiber dynamics were also studied to better understand the underlying mechanism.

Here the study is duplicated as closely as possible using the amount of information available in the published article. A 51 Node-of-Ranvier fiber was modeled using the MRG model [21]. A PSA electrode was placed over the central node of the fiber and placed in a homogeneous isotropic medium. The conductivity of the medium was set to 0.2*S/m*.The neural conduction was verified with a test action potential generated with an intracellular current clamp at the myelinated fiber’s first node of Ranvier. The block threshold was defined as the minimum HFAC peak-to-peak amplitude required to block the propagation of the test action potential and was estimated with a binary search method with a search tolerance of 1%.

The effect of the electrode-to-axon distance and the HFAC frequency is plotted in Fig 8(a) and (b) respectively. The neural dynamic of the axon’s 25^th^ node of Ranvier is shown in Fig 8(c). Measured block thresholds are very close to the one obtained in the original study from Bhadra *et al*. The dynamics of the gating particles and the membrane potential are also very similar: the membrane voltage is highly contaminated by high frequency near the electrode but less on both sides and the action potentials from the onset response are visible. As noted by Bhadra *et al*, particle values show rapid evolution and a high value of *m*, small variations of *h* around a small average value, and slow evolution of *s* and *mp* to values near 1.The evolution of block threshold with distance and frequency is also following the original study.

**Fig 8.**
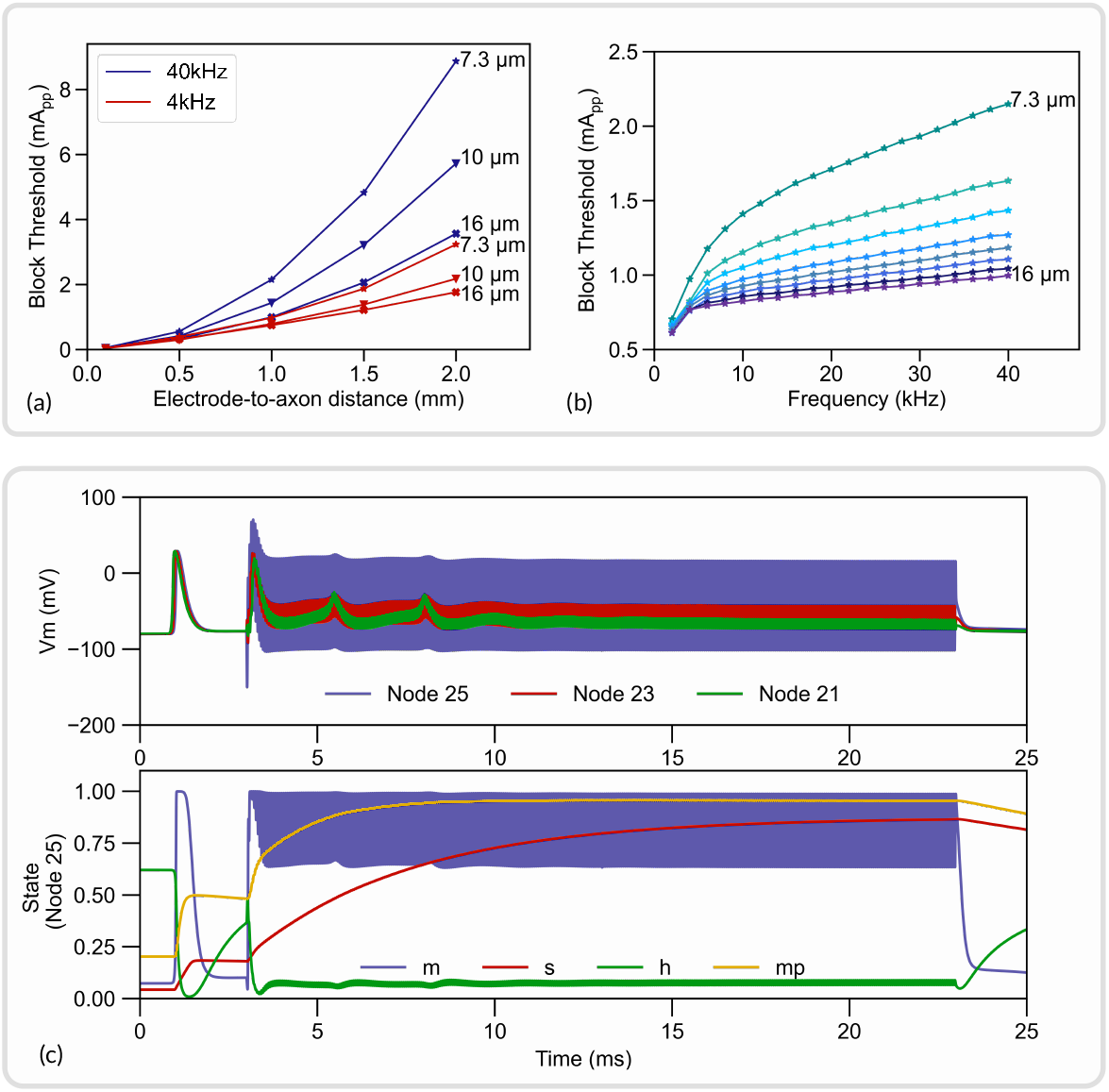
Replication of an *in-silico* study from [15] with NRV. (a) Block thresholds versus electrode-to-axon distance for a 7.3*μm*, a 10*μm* and a 16*μm* axon. Tested HFAC frequency is 4kHz and 40kHz; (b) Block thresholds versus HFAC frequency for axon diameters ranging from 7.3*μm* to 16*μm*; (c) Membrane potential (top) and gating variables (bottom) of a 10*μm* axon. An action potential is initiated at *t* = 0.5*ms* with an intracellular current clamp. A supra-blocking threshold HFAC is applied from *t* = 3*ms* to *t* = 23*ms*.

As expected, no exact match was obtained as some parameters of the simulation, such as spatial discretization, had to be guessed. Also, the study of Bhadra [15] uses the original discrete model parameters of the MRG model, whereas we use interpolated values in our framework. This first result validates the implementation of the neural models in the NRV framework. Python scripts to generate and plot this *in-silico* study are made available in the supplementary material (S3. Files). Furthermore, the iPython notebook file provided in supplementary material (S4. iPython Notebook) also demonstrated how in just a couple of lines of code this simulation can be replicated.

#### Replication of *in-vivo* studies

In this section, the *in-vivo* experiment from Nannini and Horch [64] and from Yoshida and Horch [65] were replicated with the NRV framework. Both studies aimed at characterizing the recruitment properties of LIFEs implanted in the tibial nerve of a cat. We focused our efforts on replicating the fiber recruitment versus stimulation current curve (Fig 5(a) from [64]) and the fiber recruitment versus pulse-width (Fig 2 from [65]). In both studies, fiber recruitment was estimated from recordings of the force produced by the gastrocnemius muscle (GM). As a first approximation, the cat’s tibial nerve was modeled as a mono-fascicular nerve. The fascicle innervating the GM muscle measures approximately one-fourth of the tibial nerve in diameter in the rat [66, 67] and the tibial nerve of the cat has a diameter of approximately 2.2mm [68]. Thus, the simulated GM innervating fascicle diameter was set to 550*μm*. The fascicle was filled with 500 randomly picked myelinated axons with diameters ranging from 1*μm* to 16*μm*. Axon diameter probability density was defined from fiber diameter distribution measured in the cat’s tibial nerve [69]. LIFEs were modeled following the given specification, i.e. with a diameter of 25*μm* and an active-site length of 1*mm*. Neither study provided precise information on the electrode location within the fascicle. Notably, no post-experimental histological studies were carried out around the electrode implantation zone. Consequently, *in-silico* recruitment curves were estimated with 10 different LIFEs locations randomly positioned (uniform distribution) within the fascicle.

Simulated recruitment curves with original data points superimposed are shown in Fig 9. We also define the recruitment rate as the slope of the curve evaluated between 0.2 and 0.8 of the recruitment.

**Fig 9.**
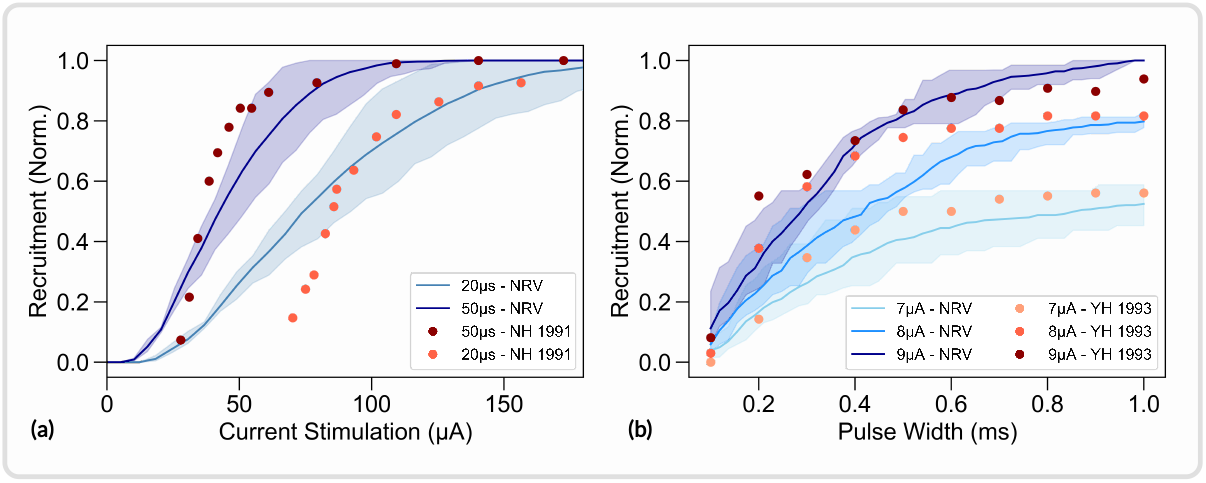
Replication of *in-vivo* studies from [64] and [65] with NRV. *In-silico* and *in-vivo* data superimposed. *In-silico* fibers are myelinated only and modelized using the MRG model [21]. (a) Recruitment versus stimulation current with a 50*μs* and 20*μs* cathodic pulse duration; (b) recruitment versus cathodic duration with a 7*μA*, 8*μA*, and 9*μA* stimulus; YH 1993 refers to data from [65]; NH 1991 refers to data from [64]. All experimental data are normalized with the maximum measured force for comparison purposes.

Simulated recruitment curves and *in-vivo* experimental data are in good agreement: reducing the pulse width from 50*μs* to 20*μs* decreases the recruitment rate by a factor of about 2.1 in the Nannima and Horch study and by about 2.4 on average in the *in-silico* study (*p <* 0.01, t-test) as shown in Fig 9(a). The stimulation amplitude does not significantly affect the recruitment rate in the simulated recruitment versus pulse width duration curves (*p >* 0.05, t-test), and the recruitment rate is increased by a factor of 1.1 only when the stimulation amplitude is increased from 7*μA* to 9*μA* in the Yoshida and Horch study (Fig 9(b)). When the stimulation pulse width is 1ms, the *in-vivo* recruitment difference between a 7*μA* and a 9*μA* pulse amplitude is about 0.37 and is about 0.48 in the *in-silico* study (*p <* 0.01, t-test). Effects of pulse duration and pulse amplitude observed in the *in-vivo* are well captured by the simulation, but the model overestimated the effect of pulse amplitude by about 22% and the effect of pulse width by about 14% when compared to the *in-vivo* data. Both *in-vivo* studies do not provide any statistical data thus the comparison is limited.

The absolute predictions of the *in-silico* recruitment curves follow reasonably closely *in-vivo* data. Dissimilarities are however observed, notably in the lower range of the stimulation current versus recruitment curves (Fig 9(a)). *In-silico* studies also underestimate the recruitment rate by about 53% on average when compared to the Nannima and Horch study (Fig 9(a)) and by about 23% on average when compared to the Yoshida and Horch study (Fig 9(b)). Such differences are to be expected due to the different nature of *in-vivo* and *in-silico* recruitment curves. *In-vivo* recruitment data are obtained from muscle force recording that is innervated by a specific and localized subset of fiber in the tibial fascicle. *In-silico* curves are obtained from the entire fiber population without spatial or functional considerations, resulting in much smoother recruitment curves.

This example also demonstrates the influence of LIFEs positioning on recruitment curves. The random positioning of LIFEs led to a standard deviation of the simulated recruitment rates of about on average 21%. This result indicates that the *in-vivo* evaluation of other stimulation parameters effect could be hidden by the variability caused by the uncontrolled positioning LIFEs during implantation. *In-silico* studies using the NRV framework facilitate the investigation of isolated stimulation parameters’ influence as each parameter can be precisely controlled and monitored. Python scripts for generating and plotting the data presented in this section are available in the supplementary material (S5. Files).

### Extracellular recordings

To demonstrate the extracellular recording capabilities of NRV, we simulated 5 monofascicular nerves comprised of 500 myelinated axons and 1059 unmyelinated axons randomly distributed according to the statistics for the sciatic nerve [69]. The corresponding radius of generated fascicles is about 120*μm*. 8 recording points are placed along the fascicle at the surface of the fascicle, each 2.5*mm*, starting at 2.5*mm* from a current clamp initiating an action potential on all fibers at the same time, as illustrated in Fig 10.

**Fig 10.**
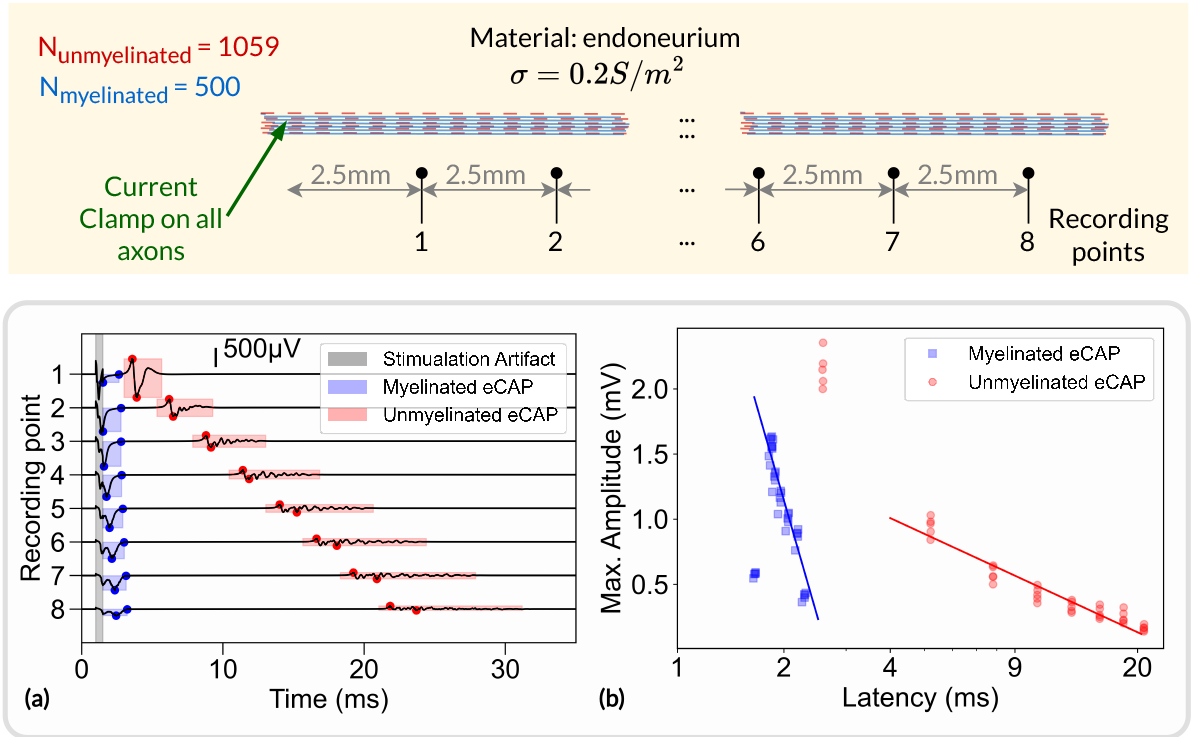
Extracellular recordings simulated with NRV. Top: schematic representation of the simulation. Myelinated and unmyelinated fibers are placed in a homogeneous infinite medium (endoneurium) and activated with a current clamp. Recording electrodes are placed along the fibers and separated by 2.5*mm*; (a) simulated potential at each recording point. Myelinated and unmyelinated eCAP contributions are highlighted in blue and red, respectively. The stimulation artifact is highlighted in gray; (b) Maximum amplitude of myelinated eCAP (in blue) and unmyelinated eCAP (in red) versus eCAP latency.

An example of evoked compound action potential (eCAP) on the 8 successive recording points is shown in Fig 10(a). Two propagating eCAPs are visible, highlighted in blue and red in the figure, and correspond to the fast propagation of action potentials in myelinated fibers and the slow propagation in unmyelinated fibers, respectively. The voltage potential resulting from the stimulation artifact is also visible. The graph shows a linear relationship between amplitude and latency (in log scale), which was also observed *in-vivo* [70]. The *in-silico* amplitude-latency slope for the myelinated fibers is about -2.0 indicating an inverse square relationship. The *in-vivo* study reported a slope of about -1.8, which is consistent with the *in-silico* estimation. It should be noted that points on the first *in-silico* recording point are not included in the fit. For myelinated eCAP, the amplitude is too small and likely obscured by the stimulation artifact. For unmyelinated eCAP, the amplitude is large, due to full synchronization of the fibers, which is unlikely to be observed in *in vivo* experiments and inherent to *in silico* simulations. Python scripts for generating and plotting the data presented in this section are available in the supplementary material (S6. Files).

### Optimizing the stimulus energy

To illustrate the optimization capabilities implemented in the NRV framework, we set up an optimization problem aimed at reducing the stimulus energy of an electrical stimulation. The problem is illustrated in Fig 11 (a). The static context of the optimization problem consists of a monofascicular nerve with a LIFE implanted in its center. The fascicle is filled with 205 myelinated axons modeled with the MRG model. The cost function of the optimization problem is defined as the sum of a stimulus energy contribution [60] and a number of recruited fibers contribution:

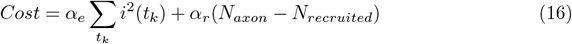

Where *t*_*k*_ is the discrete time step of the simulation, and *α*_*e*_ and *α*_*r*_ are two weighting coefficients. We chose *α*_*r*_ *>> α*_*e*_ to favor the recruitment of fiber over the stimulus energy reduction in the optimization process.

The stimulation pulse is adjusted via the NRV’s context modifier according to two scenarios:

- The stimulus is a cathodic conventional square pulse. In this scenario, both the pulse duration and pulse amplitude can be optimized, resulting in a two-dimensional optimization problem. The tuning parameters input vector *𝒳*_*sq*_ of the optimization problem is thus defined as follow:

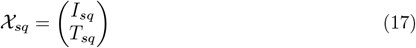

Where *I*_*sq*_ and *T*_*sq*_ are the conventional pulse amplitude and pulse width respectively.
- The stimulation is defined as an arbitrary cathodic pulse through interpolated splines over *N* points which are individually defined in time and amplitude [72]. This second optimization scenario results in a 2*N* -dimensional problem with the input vector 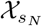 defined as follow:

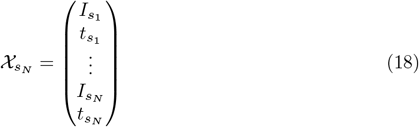

Where the pair of elements 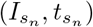 represents the simulation amplitude and time coordinates of each interpolated point.

The optimization problem was solved for the square pulse stimulus and for a 1-, 2-, and 3-points spline interpolation stimulus, resulting in four optimization problems with 2, 2, 4, and 6 dimensions respectively. A particle swarm optimization (PSO, [73]) algorithm was used as the solver, with 25 PSO particles and 60 iterations.

**Fig 11.**
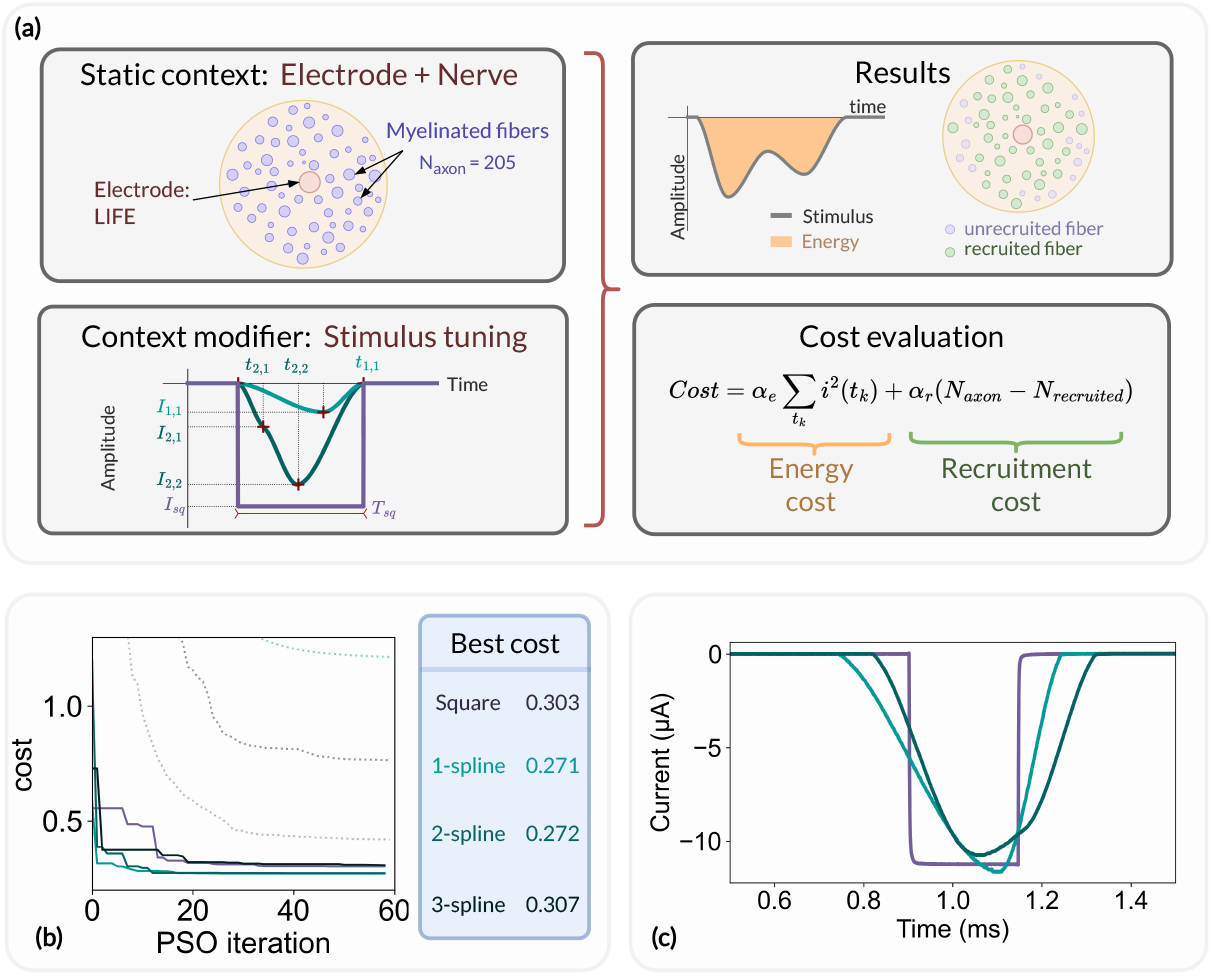
Optimization problem formulation and results. (a) Overview of the optimization problem. The simulation context consists of a monofascicular nerve and a LIFE and the stimulus parameters are modified by the context modifier after each optimization iteration (left). The simulation results are processed to derive the number of recruited fibers and the stimulus energy and then used to update the cost function (right). (b) Cost function evolution: best (solid line) and average (dotted line) cost of the swarm for a square pulse (purple), a 1- (light blue), a 2- (teal blue), and a 3- (dark blue) spline stimulus; (c) Current recording of the square pulse, 1- and 2-splines energy-optimal stimulus applied to a LIFE plunged into a saline solution using a custom arbitrary-waveform stimulator [71]. The current is recorded via a 1*k*Ω shunt resistor.

For each optimization scenario, the evolution of the best cost and average cost of the PSO particles is shown in Fig 11 (b). After the first iteration of the PSO, the average cost for each tested stimulus is greater than 1, indicating that, on average, not every fiber of the fascicle is recruited. However, at least one particle of the PSO starts with stimulus parameters that result in a cost below 1 and thus fully recruits the fascicle. In such a situation, further reduction of the cost function can only be achieved by reducing the energy of the stimulus. For square pulse stimulation, the best cost was reduced from 0.72 to 0.30 over the 60 PSO iterations, representing a cost reduction of around 40%. The square pulse energy-optimal parameters were found to be *T*_*sq*_ = 232*μs* and *I*_*sq*_ = 11*μA*. It is worth noting that the *T*_*sq*_ obtained is very close to the 200*μs* energy-optimal pulse duration found by Wongsarnpigoon and Grill [60]. The stimulus energy was further reduced with the 1-spline stimulus, resulting in an energy reduction of about 10% compared to the energy-optimal parameters of the square pulse. No significant differences were observed between the 1-spline and 2-splines stimuli, and the 3-splines stimulus showed no improvement in energy compared to the conventional square pulse.

Each aspect of the optimization problem was described using the NRV API objects, methods, and functions described in this paper. In particular, the optimization was run on multiple CPU cores using the built-in parallelization capabilities of the framework’s `Problem` and `CostFunction` classes. The parallelization resulted in a runtime for each optimization scenario of about 20 hours on a single core to about 1 hour on 50 CPU cores.

Finally, we exported the energy-optimal square pulse, 1-spline, and 2-splines stimuli to an arbitrary waveform neurostimulator developed by our group [71]. The stimuli were applied to a custom-made LIFE with similar geometric properties to those used in the static context of the optimization problem, plunged into a saline solution. The measured current for each stimulus is plotted in Fig 11 (c). The latter result demonstrates how *in-silico* results from the NRV framework can be easily translated to *in-vivo* experiments. All Python scripts and source files required to create and run the optimization problem, analyze the results, and translate the results to the neurostimulator are available in the supplementary material (S7. Files).

## Discussion

NRV is an open-source framework developed in Python that aims to improve the accessibility of *in-silico* studies for PNS stimulation. The framework uses the third-party open-source software Gmsh and FEniCS to solve the volume conduction problem and NEURON to simulate the neural dynamics of the fibers. The NRV sources and examples are freely available on GitHub (https://github.com/fkolbl/NRV). The version of the framework associated with this manuscript is *v*1.0.0 (commit n° #886: 26f02af). The framework is distributed under the CeCILL Free Software License Agreement. We hope to provide the research community with a tool for reproducible and fully open science. We also provide long-term support:

- Community for online assistance and code-sharing: https://github.com/fkolbl/NRV/discussions,
- Complete documentation and examples with granular complexity: https://nrv.readthedocs.io/en/latest/,
- Avalaibility on package manager: https://pypi.org/project/nrv-py/.

The NRV framework is designed maintained using Continuous Integration/Continuous Development workflow, and with a focus on retro-compatibility to ensure the reproducibility of scientific results.

### Framework Validation

The integration of the FEM solver into the framework was validated by comparing the results with those obtained using the well-established commercial FEM software COMSOL Multiphysics. The difference in fiber stimulation thresholds was less than 1%, validating the implantation of the FEniCS-based FEM solver. It also shows that simulation results obtained with COMSOL Multiphysics and FEniCS can be safely compared and that migration from COMSOL Multiphysics to FEniCS does not change the simulation results.

The NRV-API aims to facilitate the replication of other *in-silico* studies and was demonstrated by replicating a study of KHFAC neural conduction block by Bhadra et al. [15]. Bhadra’s results were closely replicated with NRV using only a few lines of Python code (Supplementary Material S3. Files). NRV facilitates the replication of *in-silico* studies and thus provides a tool for other researchers to build on previously published work.

Recruitment curves predicted by NRV were compared with *in-vivo* experiments published by Nannini and Horch and Yoshida and Horch [64, 65]. The *in-silico* and *in-vivo* recruitment curves were in line, and the simulation captured well the effect of pulse width and pulse amplitude on fiber recruitment profiles. The *in-silico* study also demonstrated the effect of LIFE misplacement on recruitment curves, illustrating the advantages of *in-silico* models for monitoring and assessing the influence of individual stimulation parameters. Results related to extracellular recording of neural fibers also successfully reproduced *in-vivo* observed phenomena. *In-silico* replication of *in-vivo* is also a useful approach to extend data analysis and interpretation.

Our architecture, based on a translation of physical objects or contexts (neural fibers, fascicles, and nerves, geometry, and material properties…), also enables a generic description of a simulation context. This implementation, transparent to the user, was successfully associated with optimization methods that enable the automatic exploration of parameter space and propose novel strategies.

### Comparison with other frameworks

Table 1 provides a comparison with other open-source frameworks published. From a modeling point of view, NRV makes the same simulation assumptions as most other frameworks, i.e. there is no ephaptic coupling between axons, electro-diffusion effects are neglected, and the electric field in the nerve is solved under the quasi-static hypothesis.

**Table 1.** Comparison with other published open-source frameworks.

|  | PyPNS | ASCENT | ViNERS | TxBDC | This Work |
| --- | --- | --- | --- | --- | --- |
| FEM Solver | external | external<sup>1</sup> | internal | internal | internal |
| Nerve Model | *ex-novo* | histology<br>*ex-novo* | histology<br>*ex-novo* | *ex-novo* | *ex-novo* |
| Axon Pop. | *ex-novo* | histology<br>*ex-novo* | histology<br>*ex-novo* | *ex-novo* | histology<br>*ex-novo* |
| Axon Models | HH<br>MRG | MRG<br>RA<br>Sundt<br>TGH | Sundt<br>Gaines | Passive | MRG<br>Gaines<br>HH<br>RA<br>Sundt<br>TGH |
| Electrodes | cuff | cuff<sup>2</sup> | cuff<br>array | cuff<br>intranural | PSA<br>cuff<br>LIFEs |
| Stimulus | arbitrary | arbitrary | arbitrary | mono. pulse<br>biph. pulse | arbitrary |
| Extracellular<br>Recorders | analytic<br>FEM | none | FEM | none | analytic |
| Optimization | No | No | No | No | Yes |
| Language | Python | Python | MATLAB<sup>3</sup> | Python GUI | Python |
| Commercial<br>Licensed<br>Software | COMSOL<sup>4</sup> | COMSOL<sup>4</sup> | MATLAB<sup>3</sup> | None | None |
| Multiprocessing | N/S | N/S | N/S | N/S | Yes |

Table notes: N/S not specified; HH Hodgkin-Huxley model; MRG McIntyre-Richardson-Grill model; RA Rattay-Aberham model; TGH Tigerholm model. <sup>1</sup>ASCENT automates communication between the external FEM solver (COMSOL Multiphysics) and the framework. <sup>2</sup>ASCENT provides highly realistic cuff electrode models based on commercially available products. <sup>3</sup>MATLAB, The MathWorks Inc, Massachusetts, USA. <sup>4</sup>COMSOL Multiphysics, Comsol AB, Stockholm, Sweden.

In terms of functionality, NRV compares well with other published frameworks. Commonly used myelinated and unmyelinated axon models are already implemented in the frameworks, and adding other published or custom axon models to NRV is straightforward. NRV handles multi-electrode stimulation with arbitrary waveform stimulation. Point-source electrodes are available for quick, geometry-free stimulation. Parametrized monopolar and multipolar cuff electrodes and LIFEs are implemented in the framework for more realistic simulations. Like PyPNS and ViNERS, NRV can simulate eCAP recordings. However, the recordings are calculated analytically, whereas PyPNS and ViNERS provide more accurate FEM-based eCAP simulations.

NRV is also the only published framework that integrates simulation optimization capabilities. Tuning stimulation parameters using optimization algorithms is a promising approach for improving PNS stimulation [18, 36, 60, 61, 74] and we believe that its integration into the framework is a great benefit for the end user. Also, we provide a solution where there is no restriction to which parameters can be optimized, and optimization methods can be added by contributors. Moreover, NRV was developed in close relation to arbitrary-waveform stimulation device [71]. In comparison with other approaches, we hope to provide complete solutions to explore stimulation strategies from *in-silico* to *in-vitro* or *in-vivo* experimental setup and ease the development of future electroceuticals.

### Scalability: from basic usage to cluster computing

Like ViNERS and TxDBC, NRV is self-contained, i.e. the end user does not need to use any external software during the simulation. In addition, NRV does not depend on licensed software such as MATLAB or COMSOL Multiphysics. NRV is based on Python, which is not only open-source but has grown in popularity over the last decade and is supported by a large community. Many Python packages are freely available, making the language more versatile than MATLAB. Python is also easier to deploy on any computer, cluster of computers, or supercomputer. The entire NRV framework, including third-party libraries and software, can be installed directly from the Pypi library manager. Another advantage of using the Python language is its ability to massively parallelize simulations without license restrictions. NRV automatically monitors the parallelization of simulations, which greatly reduces simulation times while being transparent to the user. Fascicle simulations are embarrassingly parallel problems, so the theoretical simulation execution time is inversely proportional to the available number of CPU cores. To validate this, we simulated 500 myelinated fibers on a computer cluster with an increasing number of allocated CPU cores and monitored the execution time (Fig. 12). The parallelization significantly reduces the simulation execution time and follows the theoretical decrease up to 22 CPU cores allocated. No significant improvement in execution time is observed beyond 22 CPU cores allocated to the framework. This is likely due to thermal throttling effects or memory bandwidth limitations. However, in the case of multiple parallelized fascicule or nerve simulations, as with meta-heuristic optimizations, one can expect further computational cost reduction with larger clusters.

**Fig 12.**
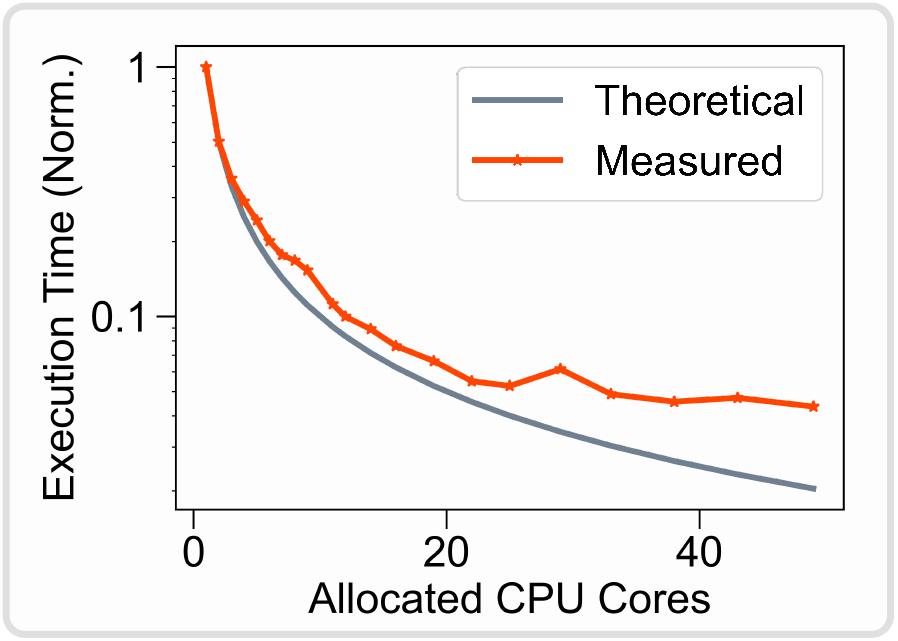
Normalized NRV simulation execution time versus allocated CPU cores count. A monofascicular nerve with 500 myelinated fibers (MRG model [21] is simulated during 5ms. Single CPU core execution time is used to normalize the data. The theoretical execution time is evaluated by dividing the single-core execution time by the number of allocated cores. Simulation is run on four Xeon Gold 6230R CPUs (104 CPU cores available) at 2.10Ghz with 1To of RAM running Ubuntu 22.04

### Perpectives

The NRV is an ongoing project that is constantly improving. To date, several improvements are under consideration. One of the limitations of the NRV framework in terms of functionality is the lack of support for histology-based realistic nerve geometry. Currently, NRV only provides classes and methods to generate parameterized cylindrical nerve and fascicle geometries, while tools to facilitate the simulation of realistic nerve geometry based on histological sections are integrated in ASCENT and ViNERS. However, no technical limitations of NRV prevent the future development of customized realistic nerves and electrodes. Since NRV is based only on open-source solutions supported by a consistent community, many different solutions and approaches can be explored to add this feature. The main challenge is to develop a solution that is easy to use for inexperienced users while being versatile enough to meet the needs of different users.

From a technical point of view, we will continue to optimize the code and extend the multiprocessing capabilities to more features of the framework. We will also explore the use of hardware acceleration, such as GPUs, to further increase computational performance. The support of interactive visualization tools, which help to validate the different steps of the simulation process as well as to facilitate data analysis, will also be considered. From a scientific point of view, we currently work on emulating complex recording methods such as Electrode Impedance Tomography [75]. Another avenue is the simulation of interconnected networks and their connection to peripheral nerves to provide *in-silico* models to more complex stimulation therapies such as vagus nerve stimulation for instance.

Updates and improvements to NRV will be continuously made available to the public on the dedicated GitHub repository. We hope that the growing number of users will increase the amount of relevant feedback that will help us to further improve the framework.

## Conclusion

In this paper, we explained the architecture, the scientific hypotheses, and the computational methods we used to implement a framework for simulating the electrical behavior of peripheral nerves. We provide a scientific tool based on fully open-source solutions. We also demonstrated the possibility of easily reproducing existing results of various kinds (*in-silico* or *in-vivo*). We also provide a solution for automated optimization of stimulation scenarios.

NRV is a versatile and powerful tool that maintains low complexity for the end user. The framework enables the simulation of a wide variety of stimulation scenarios with accurate predictions that will facilitate the development of new stimulation strategies and the transition to clinical trials. Among other things, NRV can assess the selectivity of numerous stimulation strategies but also simplifies the evaluation of more complex stimulation paradigms such as KHFAC neural conduction block. NRV also allows studies at the level of a single axon, facilitating analysis and understanding of fundamental axon behaviors. This is particularly useful when complex stimulation strategies are being studied, such as KHFAC, where the underlying mechanisms are yet not fully established [76]. NRV is also a useful tool for educational purposes and to enhance interdisciplinary research.

## Supporting information

**S1. Appendix Electrical physics validation**. Validation of the FEM equations and solver implementation on a 2-D bi-domain model.

**S2. Files Comparison between Fenics and COMSOL solvers** Python scripts to generate and plot the comparison data. Comparison data are the evaluated electrode footprint (LIFE and cuff) and the resulting activation thresholds.

**S3. Files Bhadra *et al. in-silico* study replication**. Python scripts to generate and plot the *in-silico* study replication.

**S4. iPython Notebook Bhadra *et al. in-silico* study replication with a iPython Notebook**. The iPython Notebook provides a didactic replication of the *in-silico* study.

**S5. Files Nannini and Horch, and Yoshida and Horch *in-silico* study replication data**. Python scripts to generate and plot the *in-silico* study replication.

**S6. Files Extracellular recordings *in-silico* study**. Python scripts and sources to generate and plot the *in-silico* extracellular study.

**S7. Files Energy-optimal *in-silico* study**. Python scripts and source files required to create and run the optimization problem, analyze the results, and translate the results to the neurostimulator.

## Acknowledgments

This research is part of the BioTIFS (*Improved Selectivity for Bioelectronic Therapies with Intrafascicular Stimulation*) project, and is supported by the Collaborative Research in Computational Neuroscience (CRCNS), by the the U.S. National Institutes of Health (NIH-R01-EB027584) and the French *Agence Nationale pour la Recherche* (ANR-18-NEUC0002-02).

